# Rhamnolipids from *Pseudomonas aeruginosa* Disperse the Biofilms of Sulfate-Reducing Bacteria

**DOI:** 10.1101/344150

**Authors:** Thammajun L. Wood, Lei Zhu, James Miller, Daniel S. Miller, Bei Yin, Thomas K. Wood

## Abstract

Biofilm formation is an important problem for many industries. *Desulfovibrio vulgaris* is the representative sulfate-reducing bacterium (SRB) which causes metal corrosion in oil wells and drilling equipment, and the corrosion is related to its biofilm formation. Biofilms are extremely difficult to remove since the cells are cemented in a polymer matrix. In an effort to eliminate SRB biofilms, we examined the ability of supernatants from *Pseudomonas aeruginosa* PA14 to disperse SRB biofilms. We found that the *P. aeruginosa* supernatants dispersed more than 98% of the biofilm. To determine the genetic basis of this SRB biofilm dispersal, we examined a series of *P. aeruginosa* mutants and found that mutants *rhlA*, *rhlB*, *rhlI*, and *rhlR,* defective in rhamnolipids production, had significantly reduced levels of SRB biofilm dispersal. Corroborating these results, purified rhamnolipids dispersed SRB biofilms, and rhamnolipids were detected in the *P. aeruginosa* supernatants. Hence, *P. aeruginosa* supernatants disperse SRB biofilms via rhamnolipids. In addition, the supernatants of *P. aeruginosa* dispersed the SRB biofilms more readily than protease in M9 glucose minimum medium and were also effective against biofilms of *Escherichia coli* and *Bacillus subtilis*.

## INTRODUCTION

Sulfate-reducing bacteria (SRB) are an important type of microorganisms causing iron corrosion on metal surfaces under both anaerobic and aerobic conditions^1-3^. *Desulfovibrio vulgaris* Hildenborough is a sequenced^4^ Gram-negative SRB that has been used as an SRB model organism to study biocorrosion and bioremediation of toxic metal ions^4^ as well as biofilm formation^5^, ^6^ and bioimmobilization in superfund sites^7^. It is also called the “petroleum pest” because it is commonly found in oil fields and causes “souring” of petroleum and damage to topside equipment and pipelines^8^.

Biofilms are groups of bacteria that are held together in a self-produced extracellular matrix^9^ and are difficult to remove with antimicrobial agents due to their antibiotic or biocide resistance relative to planktonic cells^10^. *Desulfovibrio* spp. populations in biofilms have a significant role for microbial induced corrosion because of their sulfide production and electron transfer mechanism^5^, and biofilms of *D. vulgaris* have been extensively shown to cause corrosion in many types of steels and other alloys^11^. The biofilms of *D. vulgaris* consists primarily of protein^5^, mannose^6^, fucose^6^, and *N*-acetylgalactosamine^6^. The biofilms of *Escherichia coli* consist primarily of proteinaceous curli fibres, flagella, and the polysaccharide cellulose^12^, and the biofilms of *Staphylococcus aureus* are largely composed of cytoplasmic proteins^13^ and extracellular genomic DNA^14^.

Many Gram-negative bacteria use quorum sensing (QS) molecules or autoinducers to communicate with each other^15^ and to form biofilms^15^. The QS mechanism can control particular processes related to cell density^16^, and QS inhibition targeting autoinducers has been used as a method to control biofilm formation^16^. The opportunistic pathogen *Pseudomonas aeruginosa* has four QS systems (Las, Rhl, Pqs, and Iqs)^17^. Each system has its own signal and regulatory protein. For the Las system, LasI synthesizes *N-*(3-oxododecanoyl)-homoserine lactone (3oxoC12HSL), and LasR is the protein receptor^17^. For the Rhl system, RhlI synthesizes *N*-butyrylhomoserine lactone, and RhlR is the protein receptor. The third QS system is quinolone-based intercellular signaling; the PQS signal is 2-heptyl-3-hydroxy-4-quinolone, and it is synthesized by the products of the PQS synthesis cluster consisting of *pqsABCD*, *phnAB*, and *pqsH*. PqsR is the protein receptor^17^. The fourth system is called Iqs, and the QS molecule is 2-(2-hydroxyphenyl)-thiazole-4-carbaldehyde^17^.

One aspect of biofilm formation controlled by the Rhl QS system is regulation of the synthesis of rhamnolipids, which are glycolipid biosurfactants composed of rhamnose and 3-(hydroxyalkanoyloxy)alkanoic acid (HAA)^18^. *P. aeruginosa* rhamnolipids affect biofilm architecture by participating in the maintenance of biofilm channels^19^ and reduce adhesion between cells^20^.

There are many reasons for forming bacterial biofilms such as a defense against stress (e.g., nutrient deprivation, antibiotics, or pH changes)^21^ and a mechanism for staying stationary in a nutrient-rich area^21^. However, bacteria also must have a means to leave the biofilm (dispersal) due to environmental changes (e.g., fluctuations in oxygen levels, alterations in nutrients, or increasing of toxic products)^22^; this process involves breaking the matrix to uncement the cells^23^.

Here, we show that supernatants from *P. aeruginosa* PA14 (henceforth PA14) disperse the biofilms of SRB, *E. coli*, and *S. aureus*. Using QS-related mutants, we determined that the biochemical basis for this dispersal is the presence of rhamnolipids in the supernatants.

## RESULTS

### PA14 wild-type supernatant disperses SRB biofilm

In an effort to investigate whether there are QS compounds utilized by the representative SRB *D. vulgaris*, we tested whether its supernatants would disperse its own biofilm formed in rich medium. *D. vulgaris* supernatant did not disperse its own biofilm within 2 h.

Since there was no negative effect of SRB supernatants on its own biofilm, we investigated the effect of the supernatant of other strains (e.g., *P. aeruginosa, P. fluorescens, Bacillus subtilis, E. coli).* Among these strains, *P. aeruginosa* PA14 and *P. aeruginosa* PAO1 dispersed SRB biofilm the most **(Fig. 1)**. The supernatants were obtained from planktonic cultures, and the SRB biofilm grown was in 96-well plates for 24 to 48 h in modified Baar’s medium. Critically, the PA14 wild-type supernatant dispersed *D. vulgaris* biofilm more than 92% after 1 to 2 h of incubation. We utilized short periods of contact of the supernatants with the SRB biofilm to avoid artifacts related with growth of the bacterium.

**Fig. 1.**
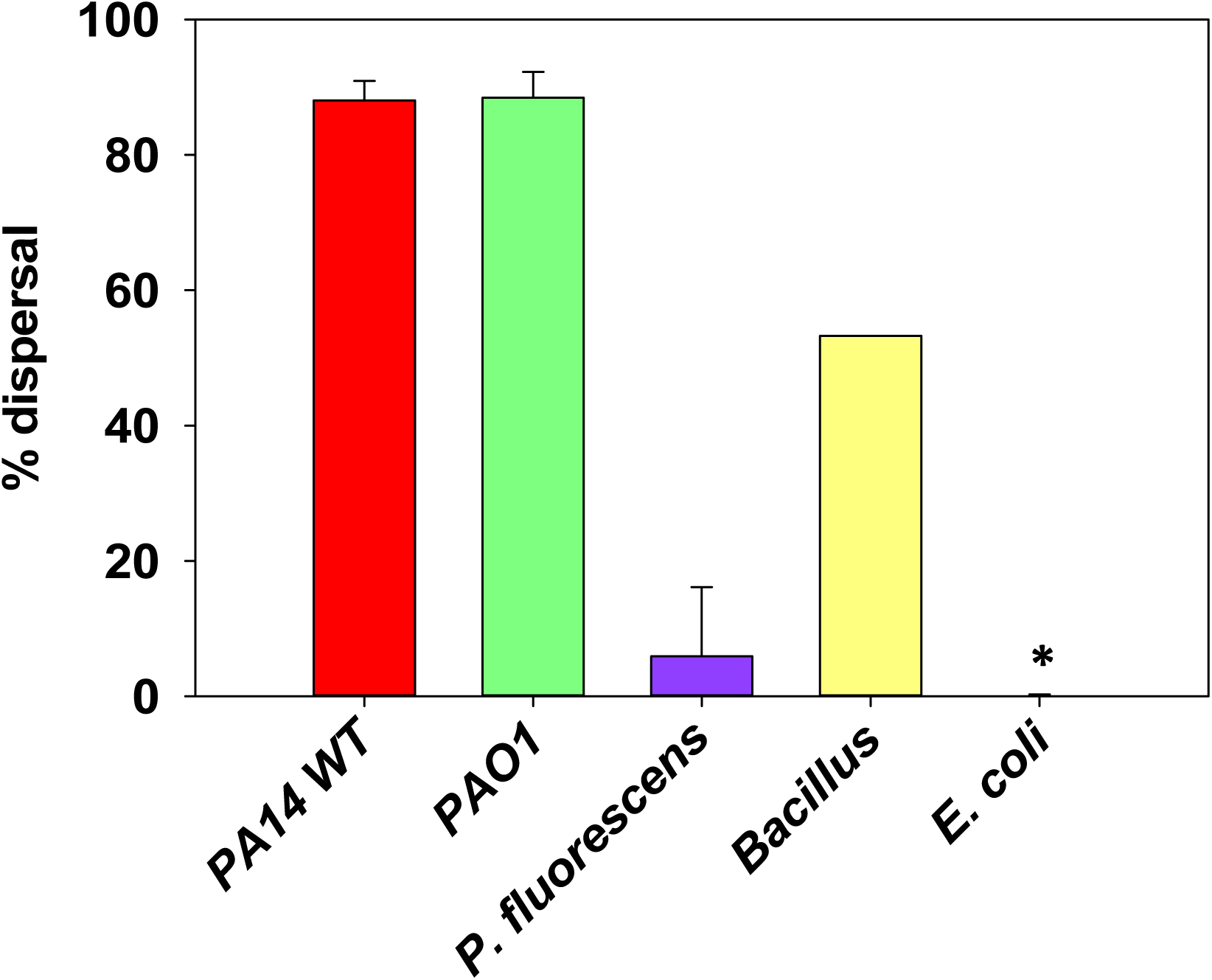
*D. vulgaris* biofilms dispersal by supernatants of *P. aeruginosa* PA14, *P. aeruginosa* PAO1, *B. subtilis, P. fluorescens,* and *E. coli* TG1. *D. vulgaris* biofilms were grown for 2 days in modified Baar’s media at 30 °C, and supernatants were concentrated to 4x and contacted with *D. vulgaris* biofilms for 2 h. * indicates no dispersal.

### Rhamnolipids in the supernatants disperse SRB biofilms

To determine mechanisms behind these strong dispersal results, we hypothesized that the compounds in the *P. aeruginosa* supernatant may be related to QS since PA14 is the best known strain for QS, and QS controls extracellular compounds like protease^24^ and rhamnolipids^24^. Hence, we tested the supernatant of mutants related to QS as well as those related to virulence factors (e.g., *lasI*^25^*, lasB*^26^*, pelA*^27^*, phzM*^28^*, phzS*^28^*, pvdF*^29^*, pchE*^30^*, lecA*^31^*, toxA*^32^*, sodM*^33^*, rhlI*^34^*, rhlR*^34^). Similar to the supernatant of the PA14 wild-type, the supernatants of the mutants *lasI* (autoinducer), *lasB* (elastase), *pelA* (oligogalacturonide lyase), *phzM* (pyocyanin)*, phzS* (pyocyanin), *pvdF* (pyoverdin), *pchE* (pyochelin), *lecA* (LecA lectin), *toxA* (exotoxinA), and *sodM* (superoxide dismutase) dispersed the *D. vulgaris* biofilm like the supernatant of PA14 wild-type **(Fig. 2),** indicating the Las QS system, Pel polysaccharide, pyocyanin, pyoverdine, pyocheline, LecA lectin, exotoxinA, superoxide dismutase had no role in the biofilm dispersal.

**Fig. 2.**
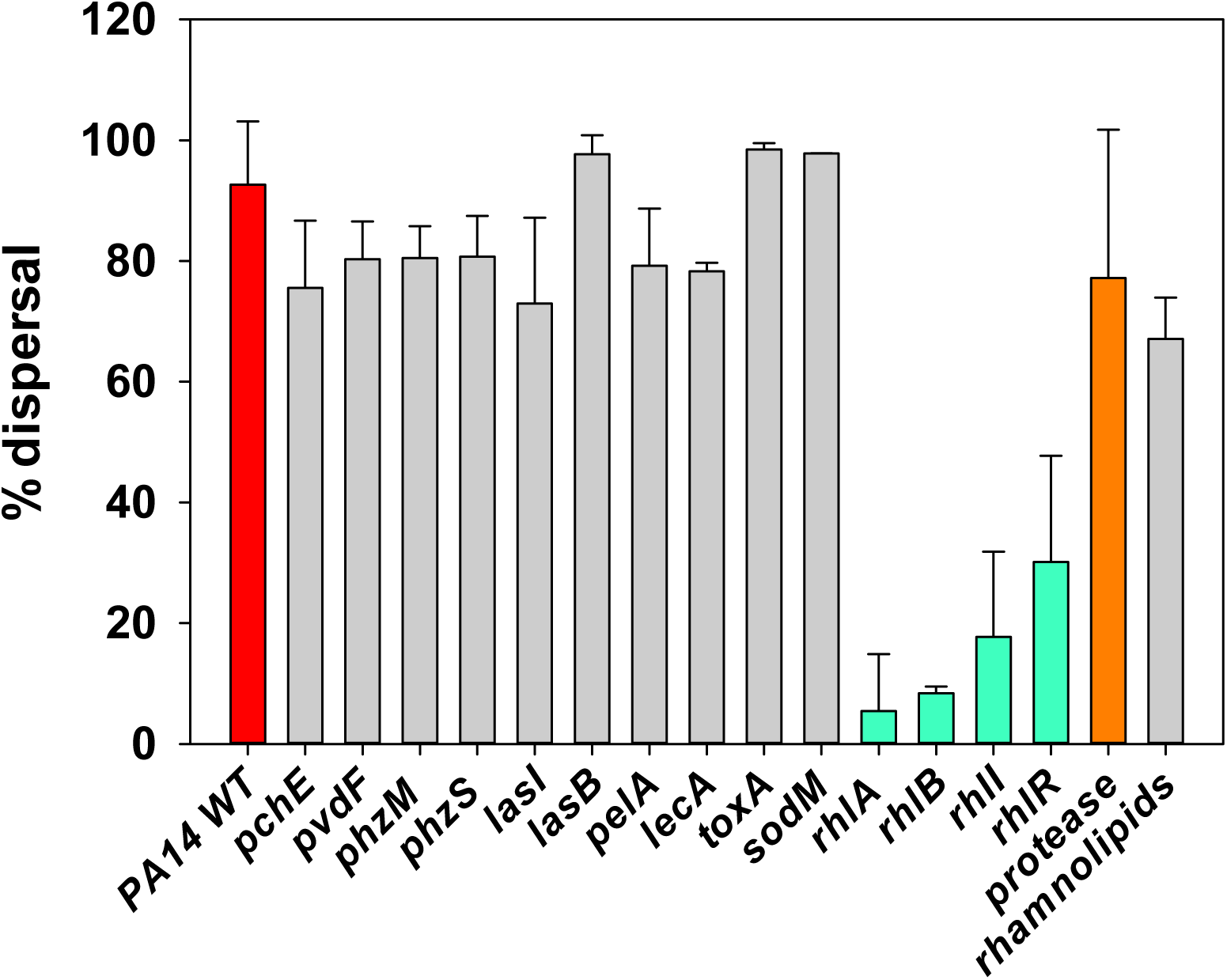
*D. vulgaris* biofilm dispersal by wild-type *P. aeruginosa* PA14 and its quorum sensing mutants. *D. vulgaris* biofilms were grown for 2 days in modified Baar’s media at 30 °C, and supernatants were concentrated to 4x and contacted with *D. vulgaris* biofilms for 2 h. Rhamnolipid standards were added at 10 mM. Protease 1 (Savinase) at 0.024 U was used as a positive control.

In contrast to the other mutants, mutations in the rhamnolipids pathway, specifically in the gene encoding the autoinducer, *rhlI,* and the gene encoding the response regulator, *rhlR*, had a pronounced decrease in dispersal (**Fig. 2**). Since the *rhlI* and *rhlR* mutants were found to affect biofilm dispersal, additional mutations in the rhamnolipid pathway were investigated: *rhlA*, which encodes rhamnosyltransferse 1 subunit A and *rhlB,* which encodes the catalytic subunit of the rhamnosyltransferase^34,35^. The supernatants of both the *rhlA* and *rhlB* mutant did not disperse SRB biofilms (**Fig. 2**). Therefore, the dispersal compounds are related to rhamnolipids.

To corroborate the 96-well biofilm dispersal results with the PA14 wild-type and *rhlB* mutant, confocal microscopy was used to visualize the remaining biofilm after treatment with supernatants from these two strains. As shown in **Fig. 3**, the supernatant from the wild-strain nearly completely dispersed the SRB biofilm whereas the supernatant from the *rhlB* mutant had no effect, just like the buffer negative control.

**Fig. 3.**
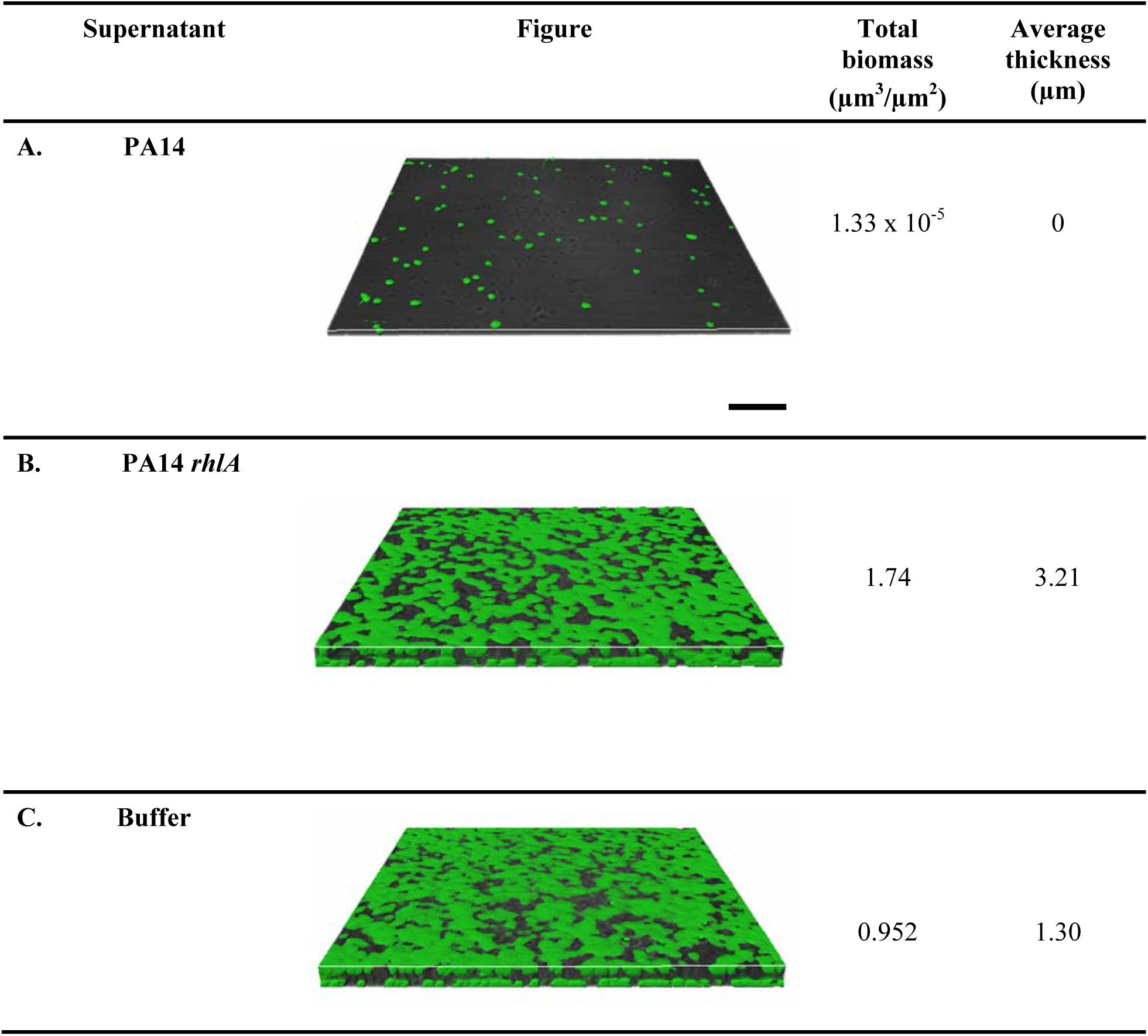
Representative images of *D. vulgaris* biofilm dispersal via *P. aeruginosa* supernatants as visualized by confocal microscopy. *D. vulgaris* biofilms were formed on an 8 chamber cell culture slide for 48 h in modified Baar’s medium, and supernatants were concentrated to 4x and contacted with *D. vulgaris* biofilms for 0.33 h. Supernatants of **(A)** *P. aeruginosa* PA14, **(B)** *P. aeruginosa* PA14 *rhlA*, and **(C)** PBS buffer. Scale bar is 50 µm.

From the rhamnolipid production pathway **(Fig. 4)**, RhlA synthesizes 3-(3-hydroxyakanoyloxy)alkanoic acids (HAA)^36^,and RhlB uses HAA to make mono-rhamnolipids^37^. Therefore, the compounds required for SRB biofilm dispersal are HAA or mono/di-rhamnolipids, respectively. However, the results from mass spectrometry showed that there was no HAA detected in the wild-type supernatant sample. Therefore we conclude that rhamnolipids are the compounds (and not HAA) that are important for biofilm dispersal.

**Fig. 4.**
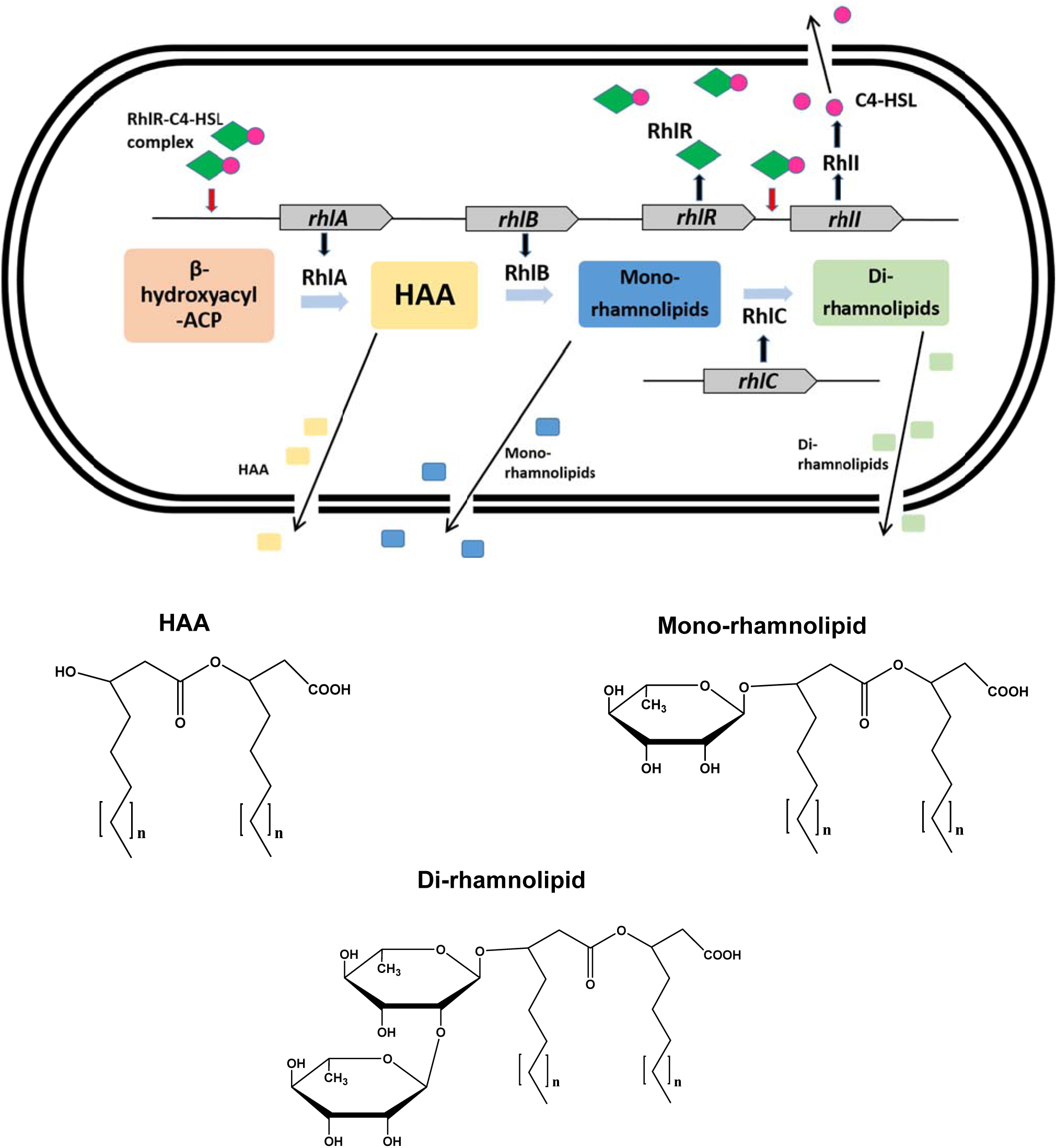
Simplified rhamnolipid biosynthesis pathway in *P. aeruginosa*^47^, ^48^. RhlR binds with *N*-butanoyl-L-homoserine lactone (C4-HSL) produced by RhlI to form a RhlR –C4-HSL complex. The RhlR –C4-HSL complex interacts with the *rhlA* promoter to initiate transcription of the *rhlAB* genes to produce rhamnolipids. HAA is 3-(hydroxyalkanoyloxy)alkanoic acid. *P. aeruginosa* commonly produces rhamnolipids containing fatty acids with chain lengths between C_8_ and C_14_ (n = 1-7)^49^.

### Rhamnolipids are present in the PA14 supernatants and disperse SRB biofilms

The genetic results clearly indicate the importance of rhamnolipids for SRB biofilm dispersal via PA14 supernatants. To corroborate these results, the supernatants of PA14, *rhlA, rhlB, rhlI,* and *rhlR* mutants were tested for the presence of rhamnolipids by mass spectrometry **(Fig. 6**). As expected, rhamnolipids were detected only in the PA14 supernatant samples and not from any of the mutant samples. There were four predominant peaks of rhamnolipids shown **(Fig. 6)** which are Rha-C_10_-C_10_, Rha-C_10_-C_12_/Rha-C_12_-C_10_, Rha-Rha-C_10_-C_10_, and Rha-Rha-C_10_-C_12/_ Rha-Rha-C_12_-C_10_. By comparing with the 10 mM rhamnolipids standard from *P. aeruginosa*, we determined the concentration of rhamnolipids in the PA14 supernatant using the total peak areas is over 100 fold less than the standard (0.1 mM, **Table 1**).

**Table 1.** Rhamnolipids in *P. aeruginosa* PA14 supernatants and in the 10 mM rhamnolipid standard. Peak areas are indicated from the mass spectrometry chromatograms.

| **Rhamnolipids<sup>51</sup>** | **Ion (m/z)** | **PA 14 WT** | **Standard** |
| --- | --- | --- | --- |
| Rha-C<sub>10</sub>-C<sub>10</sub> | 503.3 | 31 | 6374 |
| Rha-C<sub>10</sub>-C<sub>10</sub>/Rha-C<sub>12</sub>-C<sub>10</sub> | 531.3 | 3 | 294 |
| Rha-Rha-C<sub>10</sub>-C<sub>10</sub> | 649.3 | 37 | 3465 |
| Rha-C<sub>10</sub>-C<sub>12</sub>/Rha-C<sub>12</sub>-C<sub>10</sub> | 677.3 | 8 | 178 |

To show conclusively that rhamnolipids in the supernatants are the biochemical means by which the *D. vulgaris* biofilms are dispersed, purified *P. aeruginosa* rhamnolipids (10 mM) were tested. We found that the purified rhamnolipids disperse *D. vulgaris* biofilm with approximately 67% dispersal after 2 h **(Fig. 2)**. The rhamnolipids standard did not disperse the biofilm as well as the PA14 supernatant though there were more rhamnolipids in the 10 mM standard sample than the amount of rhamnolipids in the supernatant **(Table 1).** This implies that the ratio of the rhamnolipids or other compounds in the supernatant are important for SRB biofilm dispersal.

Addition of the supernatant of the *rhlB* mutant to the rhamnolipids standard did not increase the biofilm dispersal compared to the the rhamnolipids standard alone. This suggests the dispersal is due to the rhamnolipids alone and not some other product in the supernatants.

### PA14 supernatants disperse *D. desulfuricans, E. coli* MG1655, and *Staphylococcus aureus* biofilms

To test the ability of the PA14 supernatants to disperse other bacteria, biofilms of *D. desulfuricans, E. coli* MG1655, *S. aureus*, and PA14 were grown in 96-well plates, then treated with supernatants from PA14. The PA14 supernatant dispersed the biofilms of *D. desulfuricans*, *E*. *coli* MG1655, and *S. aureus* well (over 90%). Although, there are some strain specific differences, these results show that the PA14 supernatants have a great deal of utility for dispersing a wide range of biofilms.

As expected, the supernatant from the *rhlR* mutant did not disperse biofilms well (**Fig. 5**), providing further evidence of the importance of rhamnolipids. In addition, the supernatant of PA14 did not disperse its own biofilm (data not shown).

**Fig. 5.**
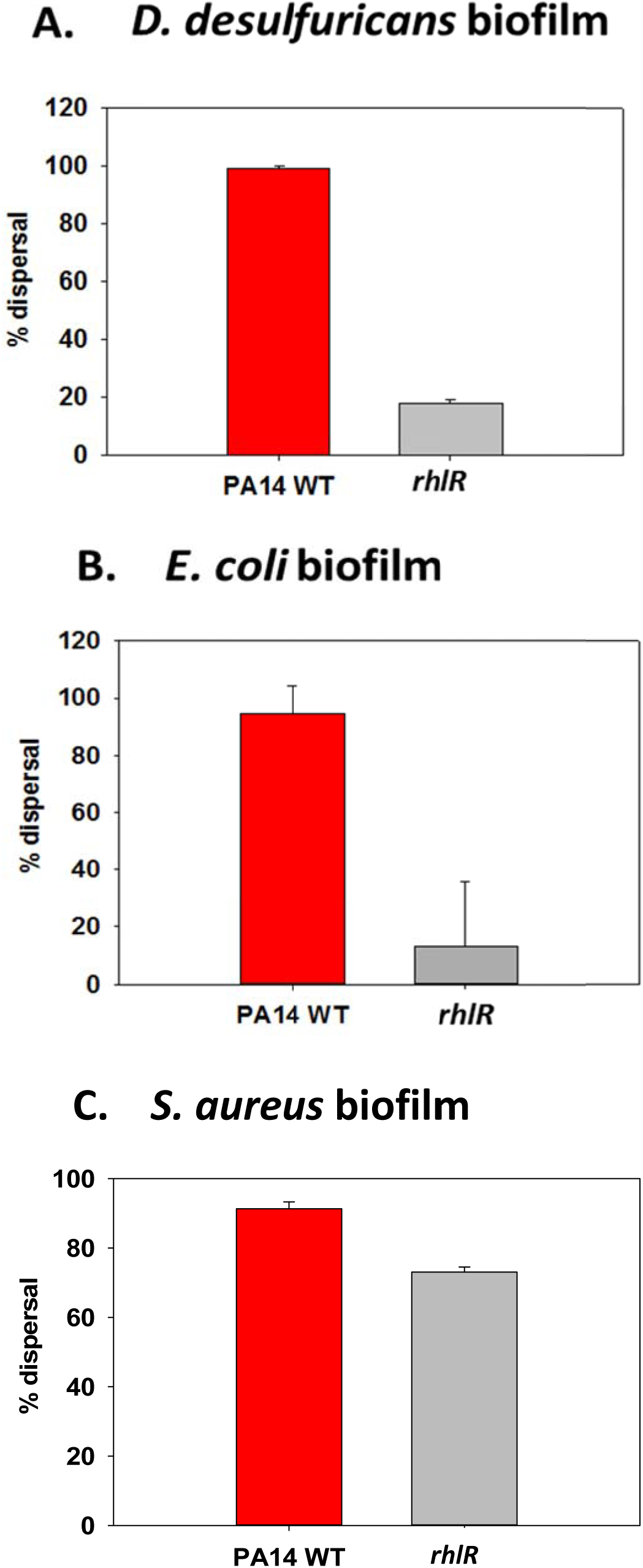
Supernatants of *P. aeruginosa* PA14 disperse biofilms of (A) *D. desulfuricans*, (B) *E. coli* MG1655 biofilm, and (C) *S. aureus*. The biofilms of *D. desulfuricans* and *E. coli* MG1655 were grown for 24 h in modified Baar’s medium and LB, respectively, and the biofilms of *S. aureus* were grown for 24 h in TSB^50^, and *P. aeruginosa* supernatants were concentrated to 4x and contacted the biofilms for 2 h except for *S. aureus* biofilm which the 1x supernatant was used instead of 4x and the contact time was 10 min.

**Fig. 6.**
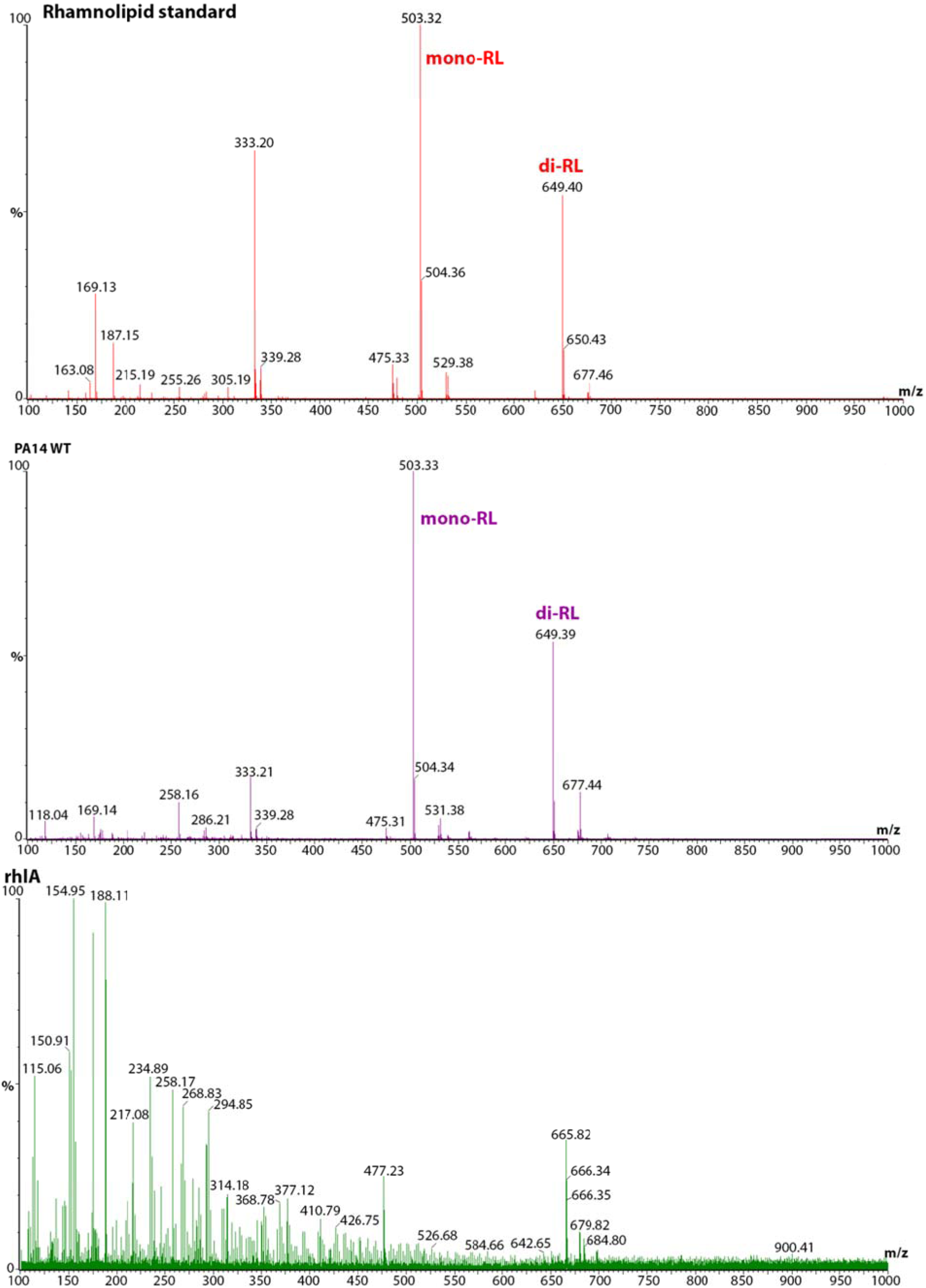
Detection of rhamnolipids in the *P. aeruginosa* PA14 supernatants via mass spectrometry. *P. aeruginosa* wild-type and the *rhlA* mutant were grown planktonically for 4 days in M9G medium at 37 °C.

### PA14 supernatant disperses SRB biofilm grown in M9G

Since biofilm matrices can change when the growth medium is altered^38^, we tested the ability of the PA14 supernatant to disperse SRB biofilms grown in minimal glucose medium (M9G). The PA14 supernatant dispersed 90-100% of the SRB biofilm when formed in modified’s Baar’s medium and dispersed about 80% of the biofilm formed in M9G. In contrast, the positive control protease, which dispersed 80% of the SRB biofilm formed in modified’s Baar’s medium, dispersed only about 40% of the biofilm grown in M9G.

## DISCUSSION

We have demonstrated that the supernatants of PA14 disperse *D. vulgaris* biofilms effectively and that the biochemical basis of this dispersal is the presence rhamnolipids. The PA14 supernatants dispersed the biofilm better than protease, and they are versatile because they also dispersed *D. desulfuricans*, *E. coli* MG1655, and *S. aureus* biofilm. This is significant since the matrix of *D. desulfuricans, E. coli*, and *S. aureus* are primarily proteins^12,13^, carbohydrates^39^, and some extracellular genomic DNA^14^, respectively, which indicates that rhamnolipids are a powerful and general approach for removing biofilms.

The rhamnolipid standard did not disperse the biofilm as well as the rhamnolipids in the PA14 WT supernatant, possibly because the ratios of the rhamnolipids are critical for biofilm dispersal activity. For example, the ratio between rhamnolipids and mono rhamnolipids affects the mixture properties (e.g., foam thickness, surface electric parameters)^40^ and changes the emulsification index and antimicrobial properties^41^. In addition, the rhamnolipid standard was not pure (90%), and the growth condition used to produce the rhamnolipids is unknown. We also surmise that purifying the rhamnolipids alters their biofilm dispersal properties. Furthermore, previously it was reported that rhamnolipids from *P. aeruginosa* together with the QS signal 3oxoC12HSL work cooperatively to disperse *E. coli* biofilms^42^; however, we found that there was no need for 3oxoC12HSL for supernatants to disperse SRB since the supernatants from the *lasI* mutant were just as effective as those from the wild-type strain (**Fig. 2**).

## METHODS

### Bacterial strains and culture conditions

*Desulfovibrio vulgaris* and *Desulfovibrio desulfuricans* were grown anaerobically in modified Baar’s medium at 30°C. PA14 wild-type and its mutants were grown in lysogeny broth (LB)^43^ or minimal medium with 0.4% glucose (M9G)^44^ at 37°C. Gentamycin (Gm) 15 μg/mL was used for culturing the PA14 mutants*. Escherichia coli* TG1, *Escherichia coli* MG1655, and *Pseudomonas fluorescens* were grown in LB at 37°C. All the strains are shown in **Table 2**. The PA14 *rhlA*, *rhlB*, *rhlI*, *rhlR*, and *lasI* mutants were verified via PCR using the primers in **Table 3**.

**Table 2.**
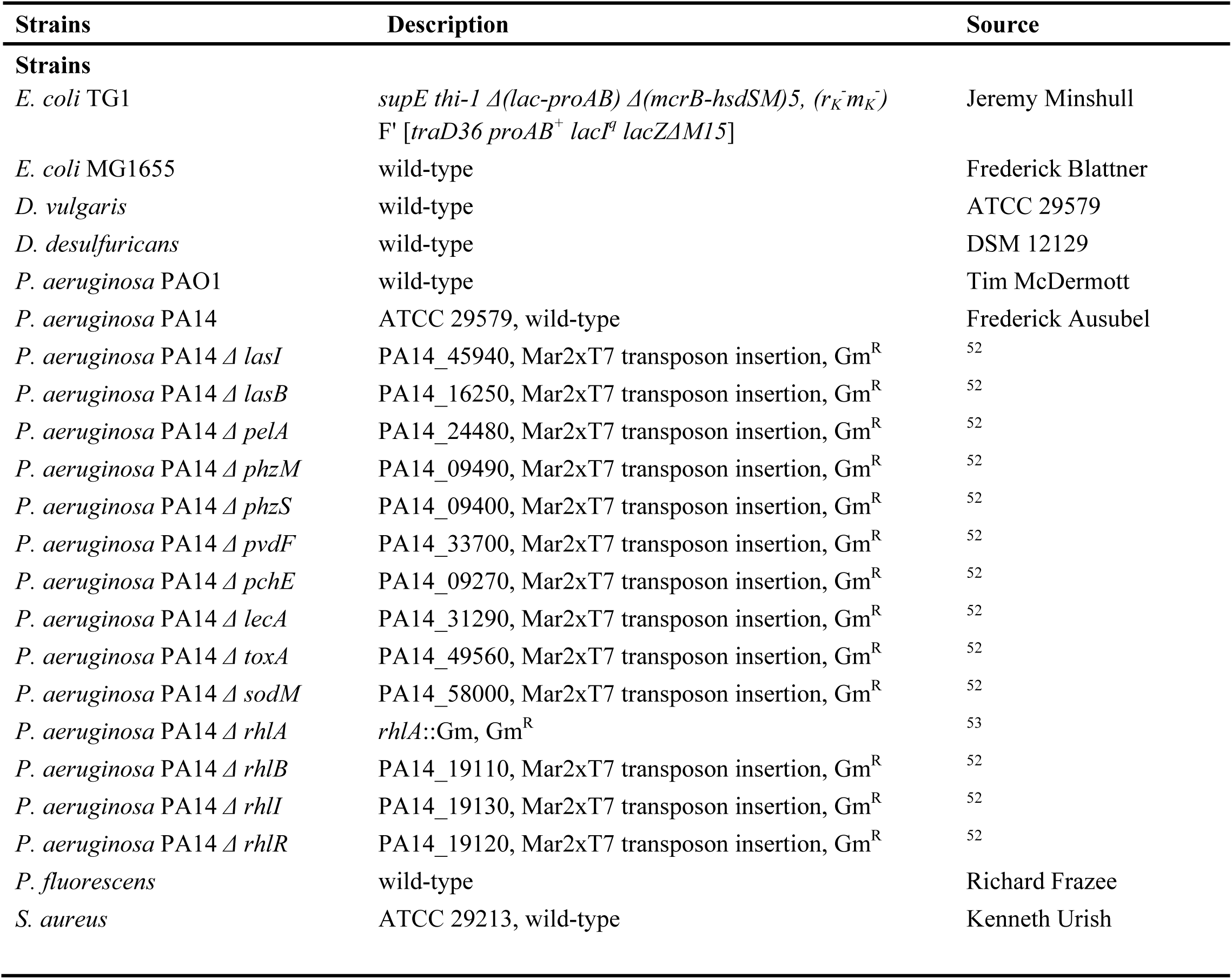
Bacterial strains used in this study.

**Table 3.**
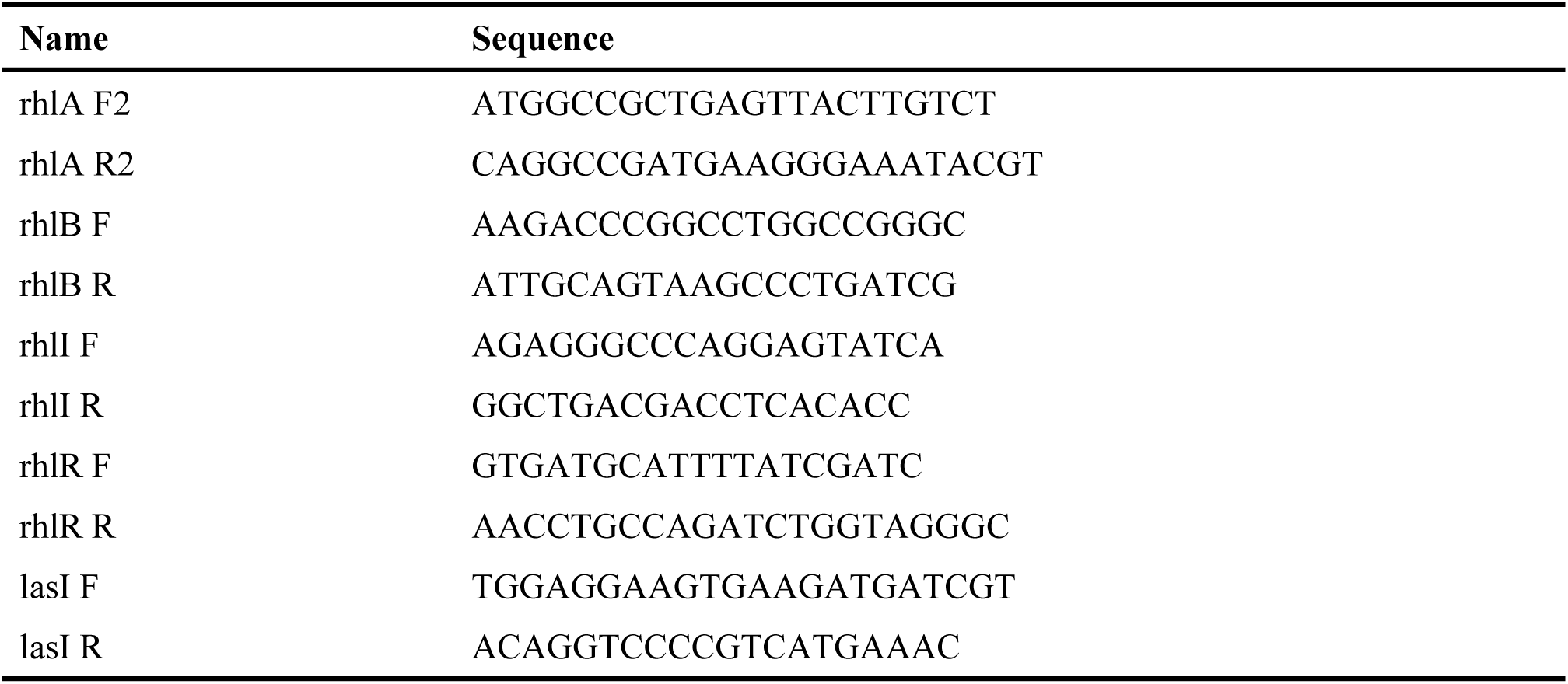
Primers used in this study. Forward primers are labeled with F, and rear primers are labeled with R.

### PA14 supernatant preparation

*P. aeruginosa* PA14 wild-type and its isogenic mutants were grown in LB or LB with Gm15 µg/mL for the mutants. The overnight cultures (200 µL) were centrifuged, the cell pellets were resuspended in 50 mL of M9G, and cultured for 3 to 4 days. The supernatants were collected and filtered using a 0.22 µm filtration system. The samples were concentrated from 2x to 4x using a speed vacuum.

### Biofilm dispersal assay using crystal violet

Biofilm formation was assayed in 96-well polystyrene plates using 0.1% crystal violet staining as described previously^45^ with some modifications. Diluted overnight cultures (150 µL) at an initial turbidity at 600 nm of 0.05 were inoculated into 96-well-plates with modified Baar’s medium, and the bacteria were cultured for 24 to 48 hours at 30°C without shaking. The planktonic cells were removed from each well, and the biofilm was washed gently using phosphate buffered saline (PBS). The supernatant or chemicals were added to the wells, and the plate was incubated at 30°C for 1 to 2 h. Then, the supernatants were removed from each well, and the remaining biofilm was washed gently using water. After the crystal violet was added to each well, the wells were rinsed and dried, and ethanol was added to dissolve the crystal violet. The total biofilm formation samples were measured at 540 nm. At least three independent cultures were used for each strain.

### Confocal Microscopy

Diluted overnight cultures (300 µL) at an initial turbidity at 600 nm of 0.1 were inoculated into an 8 chamber cell culture slide (Dot Scientific, Inc, Burton, MI) with modified Baar’s medium, and the bacteria were cultured for 48 hours at 30°C without shaking. The planktonic cells were removed from each well, and the biofilm was washed gently using PBS. Supernatants or PBS was added to the chambers, and the slide was incubated at 30°C for 20 minutes. Confocal microscopy images were taken using a PlanApo60×/1.4 oil objective lens with Olympus Fluoview 1000 confocal laser microscope. The biofilm structure images were performed using IMARIS software (Bitplane, Zurich, Switzerland), and average biomass and thickness were obtained using COMSTAT image-processing software^46^. At least three different areas were observed, and two independent cultures were tested.

### Mass spectrometric analysis of supernatants

Mass spectrometric analysis was performed on the supernatants by using a Waters Q-TOF Premier quadrupole/time-of-flight (TOF) mass spectrometer (Waters Corporation (Micromass Ltd.), Manchester, UK). Operation of the mass spectrometer was performed using MassLynx™ software Version 4.1 (http://www.waters.com). Samples were introduced into the mass spectrometer using a Waters 2695 high performance liquid chromatograph. The samples were analyzed using flow injection analysis (FIA), in which the sample is injected into the mobile phase flow and passes directly into the mass spectrometer, where the analytes are ionized and detected. The mobile phase used was 90% acetonitrile (LC-MS grade), and 10% aqueous 10 mM ammonium acetate. The flow rate was 0.15 mL/min. The nitrogen drying gas temperature was set to 300 °C at a flow of 7 L/min. The capillary voltage was 2.8 kV. The mass spectrometer was set to scan from 100-1000 m/z in negative ion mode, using electrospray ionization (ESI). Rhamnolipids standard (90%) was purchased from Sigma-Aldrich (St. Louis, MO).

## DATA AVAILABILITY

The authors declare that data supporting the findings of this study are available within the paper.

## ACKNOWLEDGMENTS

This work was supported by the Dow Chemical Company and funds derived from the Biotechnology Endowed Professorship at the Pennsylvania State University.

## AUTHOR CONTRIBUTIONS

Experiments were designed by TLW, LZ, DSM, BY, and TKW. The experiments were performed by TLW except for mass spectrometry which was performed by JM. The manuscript was written by TLW and TKW.

## COMPETING INTERESTS

The authors declare there are no competing interests.

